# Predominant myosin super-relaxed state in canine myocardium with naturally occurring dilated cardiomyopathy

**DOI:** 10.1101/2023.03.10.532046

**Authors:** Julien Ochala, Christopher T. A. Lewis, Thomas Beck, Hiroyuki Iwamoto, Anthony L. Hessel, Kenneth S. Campbell, W. Glen Pyle

## Abstract

Dilated cardiomyopathy (DCM) is a naturally occurring heart failure condition in humans and dogs, notably characterized by a reduced contractility and ejection fraction. As the identification of its underlying cellular and molecular mechanisms remain incomplete, the aim of the present study was to assess whether the molecular motor myosin and its known relaxed conformational states are altered in DCM. For that, we dissected and skinned thin cardiac strips from left ventricle obtained from six DCM Doberman Pinschers and six non-failing controls (NF). We then used a combination of Mant-ATP chase experiments and X-ray diffraction to assess both energetic and structural changes of myosin. Using the Mant-ATP chase protocol, we observed that in DCM dogs, the amount of myosin molecules in the ATP-conserving conformational state also known as super-relaxed (SRX), is significantly increased when compared with NF dogs. This alteration can be rescued by applying EMD-57033, a small molecule activating myosin. Conversely, with X-ray diffraction, we found that in DCM dogs, there is a higher proportion of myosin heads in the vicinity of actin when compared with NF dogs (1,0 to 1,1 intensity ratio). Hence, we observed an uncoupling between energetic (Mant-ATP chase) and structural (X-ray diffraction) data. Taken together, these results may indicate that in the heart of Doberman Pinschers with DCM, myosin molecules are potentially stuck in a non-sequestered but ATP-conserving SRX state, that can be counterbalanced by EMD-57033 demonstrating the potential for a myosin-centered pharmacological treatment of DCM.

**New & noteworthy:** The key finding of the present study is that, in left ventricles of dogs with a naturally occurring dilated cardiomyopathy, relaxed myosin molecules favor a non-sequestered super-relaxed state potentially impairing sarcomeric contractility. This alteration is rescuable by applying a small molecule activating myosin known as EMD-57033.

## Introduction

Dilated cardiomyopathy (DCM) is a naturally occurring disease in humans and in several large-breed dogs with Doberman Pinschers, Great Danes and Newfoundlands being at a particularly high risk. In the case of Doberman Pinschers, up to 60% of dogs are diagnosed with DCM (1, 2). The age of onset of such diseases varies but dogs usually experience cardiac dilatation and a reduced basic contractile function of the left or both ventricles (3). This diminishes ejection fraction and cardiac output, ultimately leading to ventricular wall thinning, myocardial fibrosis, congestive heart failure, and a rapid decline in the quality of life (3). Recent discoveries point to a multi-gene basis potentially involving the *PDK4* (pyruvate dehydrogenase) kinase 4 and *TTN* genes (titin) (4, 5). Despite the tremendous progress in the genetics of Doberman DCM, it remains a fatal condition in which pathophysiological causes are unclear (3, 6, 7). Thus, the precise cascade of negative cellular and molecular events triggering the diminished contractility still require exploration.

Interestingly, in Doberman Pinschers, the sensitivity to Ca^2+^ ions as well as the maximum force production of cardiac cells is dramatically impaired, indicating a potential dysfunction of myosin molecules in DCM (8). In the present study, we aimed to get insight into such molecular maladaptation by focusing on myosin relaxed states. In healthy hearts, relaxed myosin heads can either (i) interact with each other and with the filament backbone or (ii) be mobile (9). In the first case known as the super-relaxed state (SRX), the heads are sequestered, restricted from forming cross-bridges, and have a very low ATPase activity. In the second conformation called the disordered-relaxed state (DRX), myosin heads are free to bind to actin filaments and have a five to ten-fold higher ATPase activity when compared with myosin molecules in the SRX state (10, 11). Shifting from the ATP-conserving SRX to the ATP-demanding DRX state promotes a fast transition to actin binding when thin filaments are switched on by Ca^2+^ ions and is critical to cardiac performance (12). Hence, in this study, we hypothesised that in the heart of Doberman Pinschers with DCM, there is an unusually high proportion of cardiac myosin molecules blocked in the SRX conformation that (i) contributes to the observed depressed force-generating capacity and Ca^2+^ sensitivity of cardiac cells (8); and (ii) can be reversed by using a small molecule activating myosin (EMD-57033) offering an initial investigation into its therapeutic potential. To test our hypotheses, we used both non-failing controls (NF) and naturally occurring DCM dog left ventricle samples. We then isolated thin cardiac strips and performed X-ray diffraction as well as loaded Mant-ATP chase protocols to quantify structural and ATPase activity changes.

## Methods

### Canine Myocardial Samples

Animals were cared for in accordance with the guidelines of the Animal Care and Use Committee of the University of Guelph and the Canadian Council on Animal Care (8). Table 1 summarizes characteristics of NF and DCM dogs. Tissue samples were obtained from the left ventricle free wall of Doberman Pinschers with advanced DCM causing congestive heart failure (whose owners elected humane euthanasia) or dogs with no previous history of cardiovascular disease (NF). Written consent was obtained from clients for all procedures, including for the collection of myocardial samples (8). Samples were rapidly frozen in liquid nitrogen and stored at −80°C for the measurements described below.

**Table 1.** Echocardiographic measurements from Doberman Pinschers. DCM, dilated cardiomyopathy; HR, heart rate; LVIDd, left ventricle internal diameter, end diastole; LVIDs, left ventricle internal diameter, end systole; FS, fractional shortening. Non-failing group consists of 2 female and 4 male dogs. DCM group is comprised of 3 male and 3 female dogs. Data are presented as mean ± standard deviation. * denotes a significant difference when compared with NF (p < 0.05).

|  | HR (bpm) | LVIDd (mm) | LVIDs (mm) | FS (%) |
| --- | --- | --- | --- | --- |
| *NF* | 98 ± 16 | 38.7 ± 0.6 | 27.3 ± 0.8 | 29.5 ± 1.7 |
| *DCM* | 172 ± 25* | 61.3 ± 7.4* | 52.0 ± 6.6* | 15.1 ± 3.0* |

### Solutions

As previously published (13, 14), the relaxing solution contained 4 mM Mg-ATP, 1 mM free Mg^2+^, 10^-6.00^ mM free Ca^2+^, 20 mM imidazole, 7 mM EGTA, 14.5 mM creatine phosphate and KCl to adjust the ionic strength to 180 mM and pH to 7.0. Additionally, the rigor buffer for Mant-ATP chase experiments contained 120 mM K acetate, 5 mM Mg acetate, 2.5 mM K_2_HPO_4_, 50 mM MOPS, 2 mM DTT with a pH of 6.8. For experiments involving EMD-57033, all experimental solutions contained 10 μM EMD-57033 (gift from Dr Norbert Beier, E. Merck Pharmaceuticals).

### Muscle preparation and fiber permeabilization

Cryopreserved left ventricle samples were dissected into thin bundles and immersed in a membrane-permeabilising solution (relaxing solution containing glycerol; 50:50 v/v) for 24 hours at -20°C, after which they were transferred to 4°C. These bundles were kept in the membrane-permeabilising solution at 4°C for an additional 24 hours (to allow a proper skinning/membrane permeabilization process). After these steps, bundles were stored in the same buffer at -20°C for use up to one week (15, 16).

### X-ray diffraction recording and analysis

On the day of the experiments, thin cardiac strips were placed in a plastic dish containing relaxing solution. They were then transferred to the specimen chamber, filled with relaxing buffer. The ends of these thin muscle bundles were clamped at a sarcomere length of 2.00 μm. Subsequently, X-ray diffraction patterns were recorded at 15°C using a CMOS camera (Model C11440-22CU, Hamamatsu Photonics, Japan, 2048 x 2048 pixels) in combination with a 4-inch image intensifier (Model V7739PMOD, Hamamatsu Photonics, Japan). The exposure time was 500 ms. The X-ray wavelength was 0.10 nm and the specimen-to-detector distance was 2.14 m. For each preparation, approximately 20-30 diffraction patterns were recorded at the BL40XU beamline of SPring-8 and were analysed as described previously (17). To minimize radiation damage, the specimen chamber was moved by 100 μm after each exposure. Following X-ray recordings, background scattering was subtracted, and the major equatorial and myosin meridional reflections were determined as described previously (17, 18).

### Mant-ATP chase experiments

On the day of the experiments, bundles were transferred to the relaxing solution and thin cardiac strips (approximately 50 μm wide) were isolated. Their ends were individually clamped to half-split copper meshes designed for electron microscopy (SPI G100 2010C-XA, width, 3 mm), which had been glued to glass slides (Academy, 26 x 76 mm, thickness 1.00-1.20 mm). Cover slips were attached to the top using double-sided tape to create flow chambers (Menzel-Gläser, 22 x 22 mm, thickness 0.13-0.16 mm) (13, 14). Subsequently, at 25°C, for each cardiac strip, the sarcomere length was set at 2.00 μm using the brightfield mode of a Zeiss Axio Scope A1 microscope. Similar to previous studies (13, 14), each thin strip was first incubated for five minutes with a rigor buffer. A solution containing the rigor buffer with 250 μM Mant-ATP was then flushed and kept in the chamber for five minutes. At the end of this step, another solution made of the rigor buffer with 4 mM ATP was added with simultaneous acquisition of the Mant-ATP chase.

For fluorescence acquisition, a Zeiss Axio Scope A1 microscope was used with a Plan-Apochromat 20x/0.8 objective and a Zeiss AxioCam ICm 1 camera. Frames were acquired every five seconds with a 20 ms exposure time using a DAPI filter set, for 5 min. Tests were run prior to starting the current study where images were collected for 15 min instead of 5 min – these tests did not reveal any significant difference in the parameters calculated. Three regions of each individual strip were sampled for fluorescence decay using the ROI manager in ImageJ as previously published (13, 14). The mean background fluorescence intensity was subtracted from the average of the fiber fluorescence intensity (for each image taken). Each time point was then normalized by the fluorescence intensity of the final Mant-ATP image before washout (T = 0). These data were then fit to an unconstrained double exponential decay using Graphpad Prism 9.0:

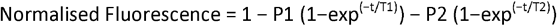

Where P1 is the amplitude of the initial rapid decay approximating the ATP-consuming DRX state with T1 as the time constant for this decay. P2 is the slower second decay approximating the proportion of myosin heads in the ATP-conserving SRX state with its associated time constant T2 (13).

### Statistical analysis

Echocardiographic data was analyzed using a one-way ANOVA. Comparison between groups of data was performed using unpaired t-tests (X-ray diffraction) or two-way ANOVA (Mant-ATP experiments, with the following two factors: NF/DCM and EMD-57033+/−). All data are expressed as means ± standard deviations; graphs were prepared and analyzed in Graphpad Prism v9. Statistical significance was set to p < 0.05.

## Results

### Higher proportion of myosin ATP-conserving SRX state in DCM dogs

To test our initial hypothesis that high number of cardiac myosin molecules from canine DCM samples were blocked in the SRX conformation, we evaluated the proportion and ATP turnover times of DRX and SRX myosin heads in isolated and permeabilized cardiac thin strips from NF and DCM dogs. For these measurements, we used a loaded Mant-ATP chase protocol to record the rate of ATP consumption of resting cardiac strips (Fig. 1A) (19). A total of 216 individual cardiac strips were tested (8 to 10 cardiac strips for each of the twelve dogs, with or without EMD-57033). DCM dogs displayed slower Mant-ATP consumption as indicated by the significantly lower P1 and higher P2 values when compared with NF dogs (Fig. 1B-C). Despite this, the T1 and T2 ATP turnover lifetimes were unchanged (Fig. 1D-E). To test the reversibility of the disrupted P1 to P2 ratio in DCM dogs, we used a commercially available pharmacological compound, EMD-57033 (EMD). EMD is a thiadiazinone derivative that is known to bind to an allosteric pocket in the myosin motor domain (20) and promote the myosin DRX state (21). In the presence of 10 μM of EMD, DCM dogs presented faster Mant-ATP consumption as shown by the significantly higher P1 and lower P2 values (Fig. 1B-C). However, again, T1 and T2 ATP turnover lifetimes remained unchanged (Fig. 1D-E).

**Figure 1.**
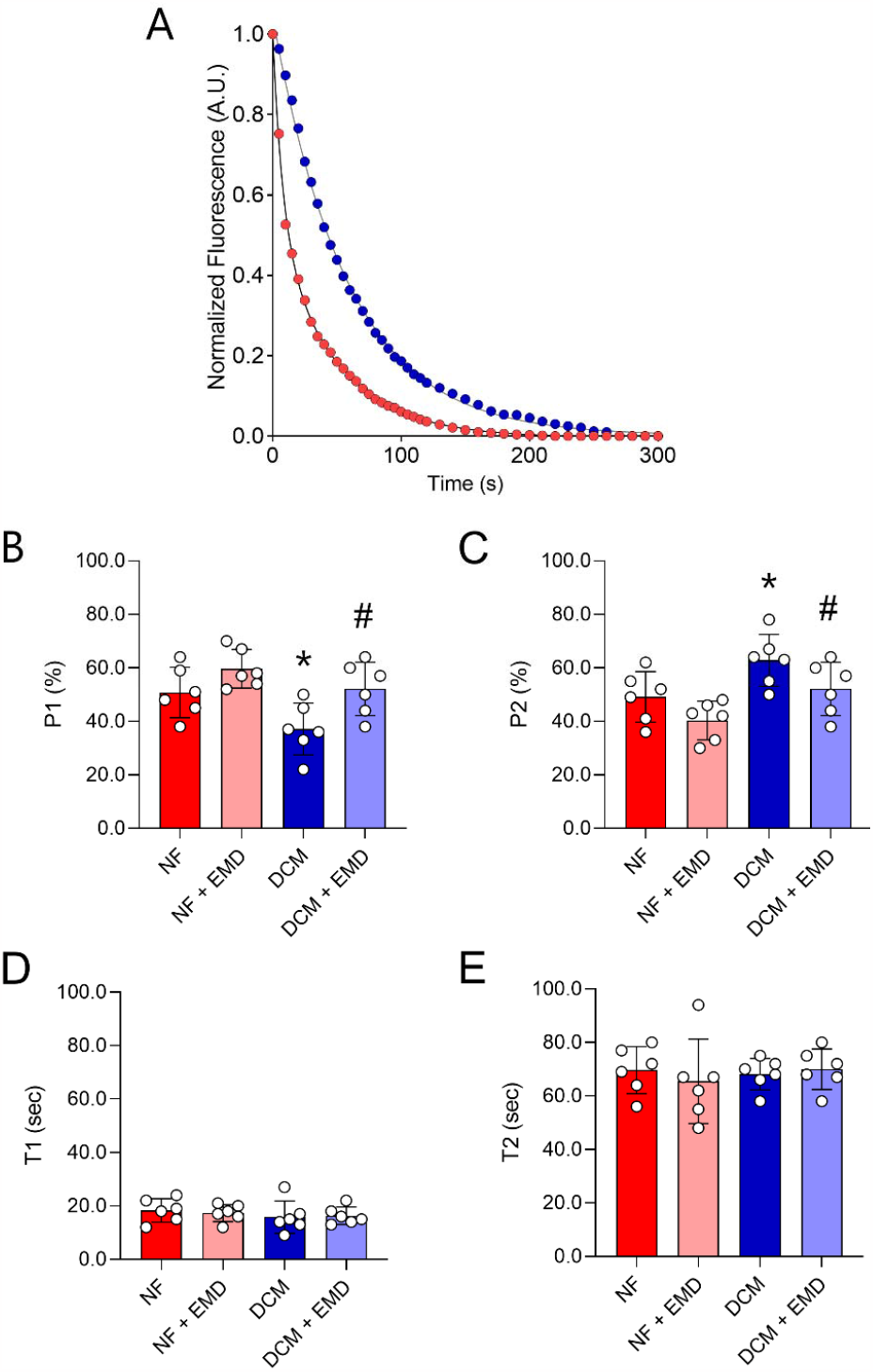
Mant-ATP data. (A) Typical Mant-ATP chase experimental data show exponential decays for thin cardiac strips isolated from NF (red) and DCM (blue) dogs. P1 (B), P2 (C), T1 (D) and T2 (E) are displayed. Dots are dog’s average data (out of 8 to 10 cardiac strips per dog). Means and standard deviations also appear on the graphs. * denotes a significant difference when compared with NF (p < 0.05). # denotes a significant difference when compared with the same group without 10 μM of EMD-57033 (EMD) (p < 0.05).

### Greater number of myosin heads in the vicinity of actin in DCM dogs

Following our finding of a greater number of myosin molecules in the SRX state in DCM dogs, we assessed whether this was accompanied by a higher proportion of myosin heads folded onto the filament backbone. For that, we assessed sarcomere structures by recording small-angle X-ray diffraction patterns and analyzed high-quality equatorial and myosin meridional reflections from isolated cardiac thin strips coming from NF and DCM dogs (Fig. 2A) (12). 1,0 and 1,1 equatorial intensities were quantified and used to generate the intensity ratio (I_1,1_ to I_1,0_) calculated, a measure of the position of myosin head mass between thick and thin filaments. Surprisingly, DCM dogs exhibited greater I_1,1_ to I_1,0_ intensity ratios values when compared with NF dogs (Fig. 2C), suggesting a mass shift towards the thin filaments, which is linked to increased cross-bridge formation upon activation (9). Despite this change, the major myosin meridional reflections, namely M3 and M6 had unchanged spacings, indicating maintained myosin crowns distances and myosin filament backbone periodicity and no expected change in cross bridge kinetics (Fig. 2C-D).

**Figure 2.**
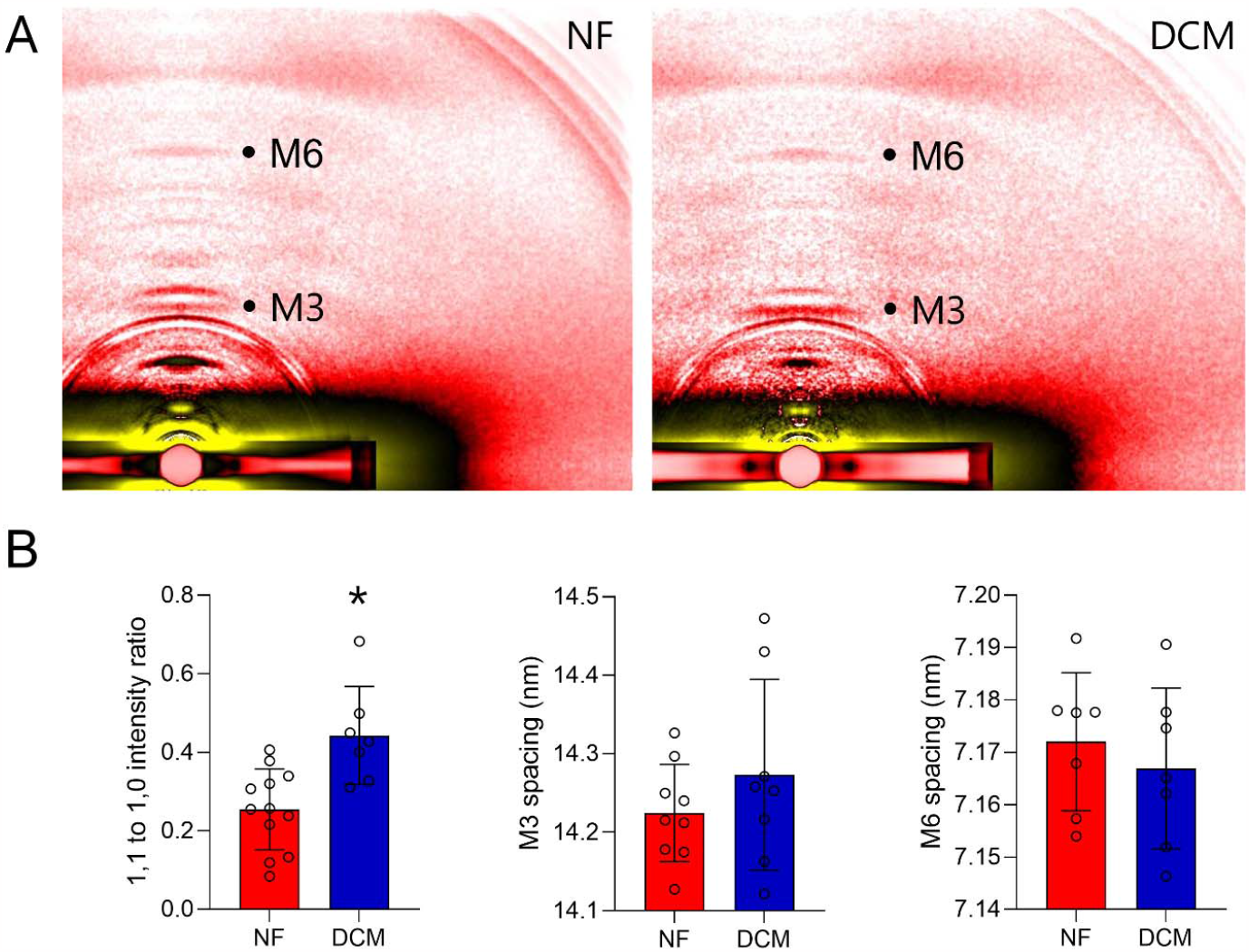
X-ray diffraction data. (A) Typical X-ray diffraction patterns obtained for thin cardiac strips isolated from NF and DCM dogs. (B) Graphs summarising the results related to the 1,1 to 1,0 intensity ratio (equatorial reflections), M3 and M6 spacing (meridional reflections). Dots are cardiac strips data. Means and standard deviations also appear on the graphs. * denotes a significant difference when compared with NF (p < 0.05).

## Discussion

In the present study, using loaded Mant-ATP chase experiments, we found that DCM dogs have an abnormally high proportion of cardiac myosin heads in the SRX state when compared with NF dogs. Using X-ray diffraction analyses, we observed that DCM dogs have an elevated level of cardiac myosin molecules positioned away from the filament backbone when compared with NF dogs, a result unexpectedly inconsistent with the SRX data. This uncoupling between energetic and structural results may indicate that in the heart of Doberman Pinschers with DCM, myosin heads are blocked in a non-sequestered but ATP-conserving SRX conformation, which can be rescued by applying a pharmacological compound (EMD) acting as a myosin activator. In agreement with our findings, recent experiments using pharmocological compounds have demostrated that the SRX state and structural myosin states can be independently changed, indicating that energetics and structural features of Ca^2+^ sensitivity are not neccesarily coupled and the underlying mechanisms for SRX to DRX and structural OFF to ON state transitions are more complex than first thought (22).

### Causes and consequences of impaired DRX to SRX ratio in DCM dogs

Myosin heads are well known to have a shutdown state in which the two heads interact with each other and with myosin sub-fragment 2 (S2) to form an interacting heads motif (IHM), leading to an ultra-low turnover of ATP (23). Even though this IHM-like *structural* state is often associated with the SRX state, it may not be necessary to get myosin molecules into the *energetic* SRX conformation (24, 25). Indeed, purified myosin sub-fragment 1 and heavy meromyosin can adopt an ATP-sparing SRX conformation on their own by stabilizing intra-molecular interactions (24, 26). Here, our observation of a mismatch between energetic (Mant-ATP chase) and structural (X-ray diffraction) data show that in the hearts of DCM dogs, the predominant SRX state may not arise from the folded-back IHM-like state but from an increase in the proportion of an intermediate configuration in which S1 and/or HMM are potentially not sequestered but have a minimal ATP usage. Studies aiming at resolving the existence of such an intermediate state are of immediate interest to the field and currently being performed.

A disrupted DRX to SRX ratio has previously been linked to the disease aetiology of human hypertrophic cardiomyopathy (HCM) due to *MYH7* or *MYPBC3* mutations (encoding for β/slow myosin heavy chain or for myosin-binding protein C, respectively) (27, 28). For instance, the *MYH7*-associated mutations cause single amino acid substitutions in the myosin head-head interface or head-tail region. These weaken the formation of the IHM crucial for myosin head sequestration (29). Hence, in HCM, the greater amount of myosin molecules in the DRX state induces an enhanced availability of myosin heads for actin binding, fitting well with the long-lasting hyper-contractility hypothesis (28, 29). Because *MYH7* or *MYPBC3* sequences are highly conserved across species, the impact of the mutations is believed to be the same among a wide range of mammal species including dogs. Human *MYH7* mutations linked to DCM appear to have totally opposite effects (29). Indeed, the variants stabilize the IHM, reducing the number of myosin heads available for recruitment. Fewer heads can enter the force-generating phase leading to hypo-contractility, with detrimental consequences for the pumping action of the heart (29, 30). In contrast to *MYH7*-associated HCM or DCM, here, the observed reduced DRX to SRX ratio cannot be caused by the direct molecular effect of the mutations on the IHM but rather to other unexplored processes impacting a non-IHM state. Cardiac myosin molecules have a turnover rate of 1-2% per day (31), they can face and be regulated by multiple post-translational modifications (32). Previously, it was shown that Doberman Pinschers with DCM have decreased phosphorylation levels of the myosin regulatory light chain (RLC) when compared with NF dogs (8). As cardiac RLC phosphorylation is thought to regulate the myosin SRX state (33, 34), we speculate that, in DCM dogs, the RLC de-phosphorylation blocks myosin heads in a non-sequestered SRX conformation, potentially impairing binding to actin and force production.

### Potential beneficial effects of EMD-57033 in DCM dogs

As Doberman Pinscher DCM is highly likely caused by multi-gene mutations (4, 5), gene therapy may not be feasible in the near future, necessitating an alternate approach in which the functional basis of the cardiac disease is targeted instead. Towards this end, small molecules directly interfering with myosin have emerged in the context of human heart failure offering great promise for veterinary medicine (35). Activating myosin pharmacologically would avoid the disadvantages of classic inotropic agents that improve cardiac contractility but also increase oxygen demand and heart rate, and are linked to hypotension, arrhythmias, and increased mortality (36). Besides Omecamtiv Mecarbil and Danicamtiv, EMD has been proven to be both a Ca^2+^-sensitizer acting on cardiac troponin and a direct myosin activator (20). Very few studies have investigated its potency in DCM. In the present study, we found a partial restoration of the DRX and SRX proportions in the presence of EMD. Even though EMD has a low bio-availability and has never been tested in clinical trials (21), our results provide evidence that other related small molecules activating myosin with similar characteristics may be potent and relevant for treating the depressed cellular force generation in DCM.

## Conclusions

Our findings indicate that, in isolated cardiac strips from DCM dogs, a non-sequestered SRX state is favored when compared with cardiac bundles from NF dogs. This adaptation can be rescued by applying myosin activators such as EMD. Altogether, our results give important new insights into the under-appreciated canine DCM pathophysiology and further strengthen the potential benefits of drugs targeting myosin activity in dogs, and also potentially in humans.

## Data availability

All the data analysed are presented in Fig. 1 and 2 but are also available upon request.

## Acknowledgement

The X-ray experiments were performed under approval of the SPring-8 Proposal Review Committee (2022A1069). X-ray data reduction and analysis performed by Accelerated Muscle Biotechnologies Consultants LLC (USA). EMD-57033 was a gift from Dr. Norbert Beier from Merck KGaA, Darmstadt, Germany. Finally, we are grateful to Prof. Thomas C. Irving for his critical review of this paper.

## Author Contributions

JO, KSC and WGP conceived the study; JO and WGP acquired funding; JO and WGP managed the project; JO, CTAL, TB, HI, ALH, KSC and WGP performed experiments; JO, CTAL, TB, HI, ALH, KSC and WGP analyzed data and interpreted the results; JO, CTAL, TB, HI, ALH, KSC and WGP wrote and reviewed the manuscript.

## References

1. Hazlett MJ, Maxie MG, Allen DG, and Wilcock BP. A retrospective study of heart disease in doberman pinscher dogs. Can Vet J 24: 205–210, 1983.

2. Wess G, Schulze A, Butz V, Simak J, Killich M, Keller LJ, Maeurer J, and Hartmann K. Prevalence of dilated cardiomyopathy in Doberman Pinschers in various age groups. J Vet Intern Med 24: 533–538, 2010.

3. O’Grady MR, Minors SL, O’Sullivan ML, and Horne R. Effect of pimobendan on case fatality rate in Doberman Pinschers with congestive heart failure caused by dilated cardiomyopathy. J Vet Intern Med 22: 897–904, 2008.

4. Meurs KM, Stern JA, Adin D, Keene BW, De Francesco TC, and Tou SP. Assessment of PDK4 and TTN gene variants in 48 Doberman Pinschers with dilated cardiomyopathy. J Am Vet Med Assoc 257: 1041–1044, 2020.

5. Simpson S, Edwards J, Emes RD, Cobb MA, Mongan NP, and Rutland CS. A predictive model for canine dilated cardiomyopathy-a meta-analysis of Doberman Pinscher data. PeerJ 3: e842, 2015.

6. Summerfield NJ, Boswood A, O’Grady MR, Gordon SG, Dukes-McEwan J, Oyama MA, Smith S, Patteson M, French AT, Culshaw GJ, Braz-Ruivo L, Estrada A, O’Sullivan ML, Loureiro J, Willis R, and Watson P. Efficacy of pimobendan in the prevention of congestive heart failure or sudden death in Doberman Pinschers with preclinical dilated cardiomyopathy (the PROTECT Study). J Vet Intern Med 26: 1337–1349, 2012.

7. O’Sullivan ML, O’Grady MR, Pyle WG, and Dawson JF. Evaluation of 10 genes encoding cardiac proteins in Doberman Pinschers with dilated cardiomyopathy. Am J Vet Res 72: 932–939, 2011.

8. Cheng Y, Hogarth KA, O’Sullivan ML, Regnier M, and Pyle WG. 2-Deoxyadenosine triphosphate restores the contractile function of cardiac myofibril from adult dogs with naturally occurring dilated cardiomyopathy. Am J Physiol Heart Circ Physiol 310: H80–91, 2016.

9. Ma W, and Irving TC. Small Angle X-ray Diffraction as a Tool for Structural Characterization of Muscle Disease. Int J Mol Sci 23: 2022.

10. McNamara JW, Li A, Dos Remedios CG, and Cooke R. The role of super-relaxed myosin in skeletal and cardiac muscle. Biophys Rev 7: 5–14, 2015.

11. Lewis CTA, and Ochala J. Myosin Heavy Chain as a Novel Key Modulator of Striated Muscle Resting State. Physiology (Bethesda) 38: 0, 2023.

12. Ma W, Henze M, Anderson RL, Gong H, Wong FL, Del Rio CL, and Irving T. The Super-Relaxed State and Length Dependent Activation in Porcine Myocardium. Circ Res 129: 617–630, 2021.

13. Ochala J, Finno CJ, and Valberg SJ. Myofibre Hyper-Contractility in Horses Expressing the Myosin Heavy Chain Myopathy Mutation, MYH1(E321G). Cells 10: 2021.

14. Ranu N, Laitila J, Dugdale HF, Mariano J, Kolb JS, Wallgren-Pettersson C, Witting N, Vissing J, Vilchez JJ, Fiorillo C, Zanoteli E, Auranen M, Jokela M, Tasca G, Claeys KG, Voermans NC, Palmio J, Huovinen S, Moggio M, Beck TN, Kontrogianni-Konstantopoulos A, Granzier H, and Ochala J. NEB mutations disrupt the super-relaxed state of myosin and remodel the muscle metabolic proteome in nemaline myopathy. Acta Neuropathol Commun 10: 185, 2022.

15. Ross JA, Levy Y, Ripolone M, Kolb JS, Turmaine M, Holt M, Lindqvist J, Claeys KG, Weis J, Monforte M, Tasca G, Moggio M, Figeac N, Zammit PS, Jungbluth H, Fiorillo C, Vissing J, Witting N, Granzier H, Zanoteli E, Hardeman EC, Wallgren-Pettersson C, and Ochala J. Impairments in contractility and cytoskeletal organisation cause nuclear defects in nemaline myopathy. Acta Neuropathol 138: 477–495, 2019.

16. Ross JA, Tasfaout H, Levy Y, Morgan J, Cowling BS, Laporte J, Zanoteli E, Romero NB, Lowe DA, Jungbluth H, Lawlor MW, Mack DL, and Ochala J. rAAV-related therapy fully rescues myonuclear and myofilament function in X-linked myotubular myopathy. Acta Neuropathol Commun 8: 167, 2020.

17. Ochala J, Iwamoto H, Larsson L, and Yagi N. A myopathy-linked tropomyosin mutation severely alters thin filament conformational changes during activation. Proc Natl Acad Sci U S A 107: 9807–9812, 2010.

18. Li M, Ogilvie H, Ochala J, Artemenko K, Iwamoto H, Yagi N, Bergquist J, and Larsson L. Aberrant post-translational modifications compromise human myosin motor function in old age. Aging Cell 14: 228–235, 2015.

19. Hooijman P, Stewart MA, and Cooke R. A new state of cardiac myosin with very slow ATP turnover: a potential cardioprotective mechanism in the heart. Biophys J 100: 1969–1976, 2011.

20. Radke MB, Taft MH, Stapel B, Hilfiker-Kleiner D, Preller M, and Manstein DJ. Small molecule-mediated refolding and activation of myosin motor function. Elife 3: e01603, 2014.

21. Jani V, Qian W, Yuan S, Irving T, and Ma W. EMD-57033 Augments the Contractility in Porcine Myocardium by Promoting the Activation of Myosin in Thick Filaments. Int J Mol Sci 23: 2022.

22. Ma W, McMillen TS, Childers MC, Gong H, Regnier M, and Irving T. Structural OFF/ON transitions of myosin in relaxed porcine myocardium predict calcium-activated force. Proc Natl Acad Sci U S A 120: e2207615120, 2023.

23. Woodhead JL, Zhao FQ, Craig R, Egelman EH, Alamo L, and Padron R. Atomic model of a myosin filament in the relaxed state. Nature 436: 1195–1199, 2005.

24. Rohde JA, Roopnarine O, Thomas DD, and Muretta JM. Mavacamten stabilizes an autoinhibited state of two-headed cardiac myosin. Proc Natl Acad Sci U S A 115: E7486–E7494, 2018.

25. Nag S, and Trivedi DV. To lie or not to lie: Super-relaxing with myosins. Elife 10: 2021.

26. Anderson RL, Trivedi DV, Sarkar SS, Henze M, Ma W, Gong H, Rogers CS, Gorham JM, Wong FL, Morck MM, Seidman JG, Ruppel KM, Irving TC, Cooke R, Green EM, and Spudich JA. Deciphering the super relaxed state of human beta-cardiac myosin and the mode of action of mavacamten from myosin molecules to muscle fibers. Proc Natl Acad Sci U S A 115: E8143–E8152, 2018.

27. Toepfer CN, Garfinkel AC, Venturini G, Wakimoto H, Repetti G, Alamo L, Sharma A, Agarwal R, Ewoldt JF, Cloonan P, Letendre J, Lun M, Olivotto I, Colan S, Ashley E, Jacoby D, Michels M, Redwood CS, Watkins HC, Day SM, Staples JF, Padron R, Chopra A, Ho CY, Chen CS, Pereira AC, Seidman JG, and Seidman CE. Myosin Sequestration Regulates Sarcomere Function, Cardiomyocyte Energetics, and Metabolism, Informing the Pathogenesis of Hypertrophic Cardiomyopathy. Circulation 141: 828–842, 2020.

28. Toepfer CN, Wakimoto H, Garfinkel AC, McDonough B, Liao D, Jiang J, Tai AC, Gorham JM, Lunde IG, Lun M, Lynch TLt, McNamara JW, Sadayappan S, Redwood CS, Watkins HC, Seidman JG, and Seidman CE. Hypertrophic cardiomyopathy mutations in MYBPC3 dysregulate myosin. Sci Transl Med 11: 2019.

29. Alamo L, Ware JS, Pinto A, Gillilan RE, Seidman JG, Seidman CE, and Padron R. Effects of myosin variants on interacting-heads motif explain distinct hypertrophic and dilated cardiomyopathy phenotypes. Elife 6: 2017.

30. Rasicci DV, Tiwari P, Bodt SML, Desetty R, Sadler FR, Sivaramakrishnan S, Craig R, and Yengo CM. Dilated cardiomyopathy mutation E525K in human beta-cardiac myosin stabilizes the interacting-heads motif and super-relaxed state of myosin. Elife 11: 2022.

31. Smith K, and Rennie MJ. The measurement of tissue protein turnover. Baillieres Clin Endocrinol Metab 10: 469–495, 1996.

32. Vanhooren V, Navarrete Santos A, Voutetakis K, Petropoulos I, Libert C, Simm A, Gonos ES, and Friguet B. Protein modification and maintenance systems as biomarkers of ageing. Mech Ageing Dev 151: 71–84, 2015.

33. Kampourakis T, and Irving M. Phosphorylation of myosin regulatory light chain controls myosin head conformation in cardiac muscle. J Mol Cell Cardiol 85: 199–206, 2015.

34. Scruggs SB, Hinken AC, Thawornkaiwong A, Robbins J, Walker LA, de Tombe PP, Geenen DL, Buttrick PM, and Solaro RJ. Ablation of ventricular myosin regulatory light chain phosphorylation in mice causes cardiac dysfunction in situ and affects neighboring myofilament protein phosphorylation. J Biol Chem 284: 5097–5106, 2009.

35. Lehman SJ, Crocini C, and Leinwand LA. Targeting the sarcomere in inherited cardiomyopathies. Nat Rev Cardiol 19: 353–363, 2022.

36. Alsulami K, and Marston S. Small Molecules acting on Myofilaments as Treatments for Heart and Skeletal Muscle Diseases. Int J Mol Sci 21: 2020.

